# The initiation of the wound healing program is regulated by the convergence of mechanical and epigenetic cues

**DOI:** 10.1101/2021.10.09.463764

**Authors:** Tanay Bhatt, Rakesh Dey, Akshay M. Hegde, Alhad Ashok Ketkar, Ajai J. Pulianmackal, Ashim P. Deb, Shravanti Rampalli, Colin Jamora

## Abstract

Wound healing in the skin is a complex physiological process that is a product of a cell state transition from homeostasis to repair. Mechanical cues are increasingly being recognized as important regulators of cellular reprogramming, but the mechanism by which it is translated to changes in gene expression and ultimately cellular behavior remains largely a mystery. To probe the molecular underpinnings of this phenomenon further, we used the downregulation of caspase-8 as a biomarker of a cell entering the wound-healing program. We found that the wound-induced release of tension within the epidermis leads to the alteration of gene expression via the nuclear translocation of the DNA methyltransferase 3A (DNMT3a). This enzyme then methylates promoters of genes that are known to be downregulated in response to wound stimuli as well as potentially novel players in the repair program. Overall, these findings illuminate the convergence of mechanical and epigenetic signaling modules that are important regulators of the transcriptome landscape required to initiate the tissue repair process in the differentiated layers of the epidermis.

**Graphical Abstract:** 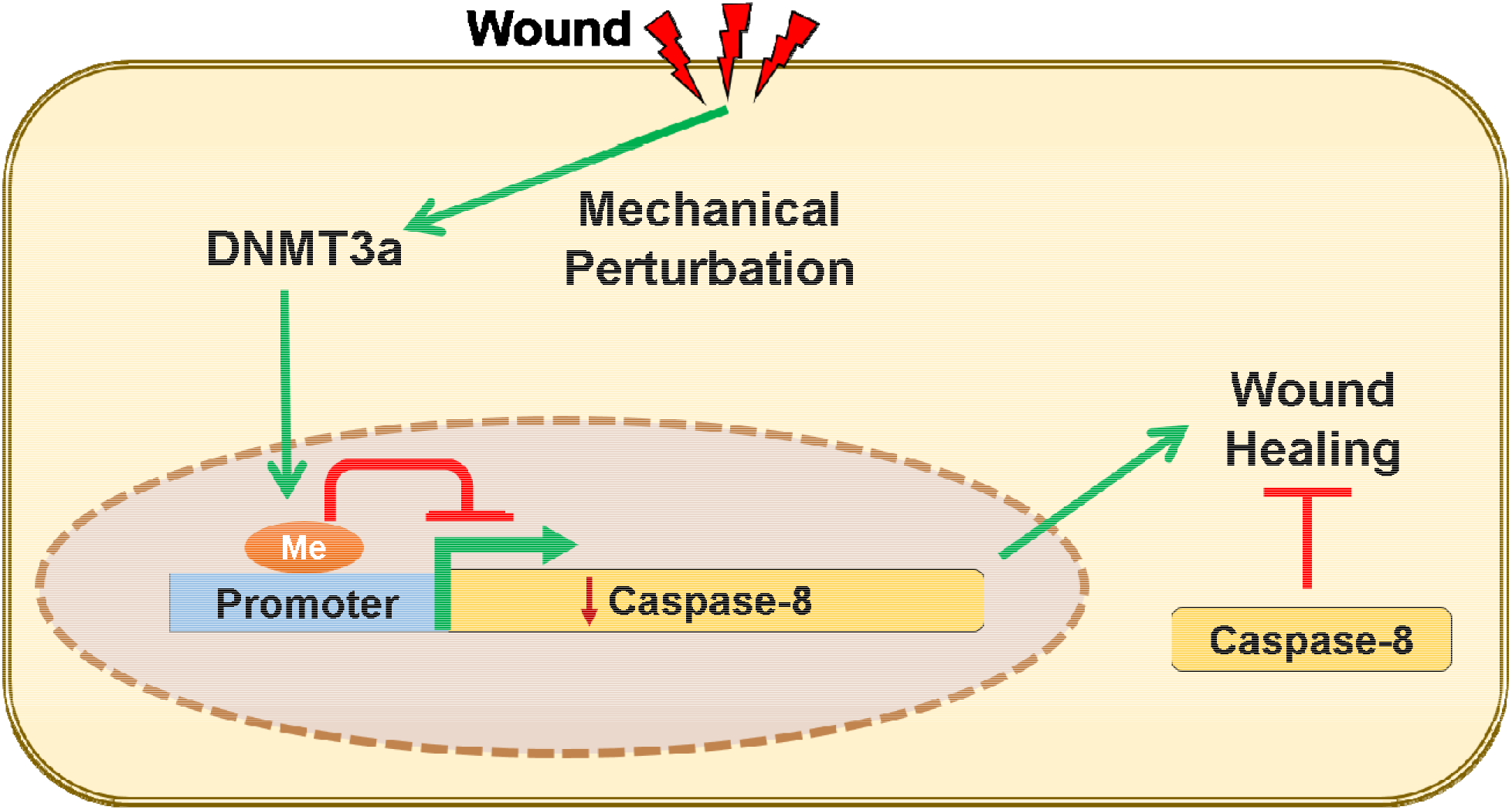

## Introduction

The wound healing program in an epithelial tissue is fundamentally a product of cell state transitions from homeostasis to a repair program. In particular, cutaneous wound healing in the adult is an intricately regulated system wherein keratinocytes and many other cell lineages exhibit their plasticity as they undergo reprogramming, to carry out otherwise dormant functions, to rebuild the damaged skin. Many of the phenomena that occur in the repair process in adult skin are, in fact, reminiscent of cellular events that operate during fetal development (1). At the other extreme, inappropriate activation of these repair processes can manifest as tissue pathology, which forms the foundation of the perception of diseases with a “wound signature” (2). The question that arises is how the whole scale changes in gene expression are accomplished in order to facilitate this cellular reprogramming.

Recently, epigenetic regulators have emerged as a vital component capable of transiently rewiring the cell’s transcriptional program to mediate the continual regeneration of the mouse epidermis (3, 4). This mode of gene regulation operates at multiple levels ranging from histone and DNA modifications, chromatin remodeling, and activity of various subtypes of RNA species such as non-coding RNAs and micro-RNAs (miRNAs) (5, 6). These epigenetic mechanisms can thus have a profound impact on the transcriptional landscape of the cell, and can easily be envisioned to participate in the transient activation or repression of the ∼1000 genes that are required for wound closure (7). However, relative to the appreciation of epigenetics in epidermal homeostasis, the understanding of its role in wound healing remains an area ripe for further exploration. Circumstantial evidence in support of a role for epigenetics in tissue repair comes from reports of their dynamic expression following injury to the skin. For instance, Ezh2, Suz12, and Eed, which are components of the Polycomb Repressive Complex 2 (PRC2), are downregulated, whereas the histone methylases JMJD3 and Utx are upregulated upon tissue damage and all return to homeostatic levels upon the completion of wound closure (8). While the description of various epigenetic players in epidermal homeostasis and wound-healing are reported, the identity and function of their upstream regulators are, to a large extent, absent in the literature.

An intriguing candidate for an upstream regulator in a highly tensile tissue such as the epidermis, are mechanical cues. The epidermis is a stratified epithelium comprised of a basal layer of proliferation competent keratincytes and suprabasal layers of differentiated cells glued together via intercellular adhesion complexes that partly endows the tissue with its barrier function. In different cell types, changes in mechanical tension have been documented to induce the nuclear translocation of important transcription factors - a notable example of which is YAP/TAZ that has proliferation stimulating gene targets (9). Many studies, including those on epidermal homeostasis and wound healing, have primarily focused on the changes in gene expression in proliferating cells (10, 11). On the other hand, differentiated cells, such as the suprabasal keratinocytes near the surface of the epidermis, have largely been relegated to bystander status. In spite of this, a few reports suggest that these neglected pools of differentiated cells are not inert in the cellular crosstalk that mediates the early responses of the tissue to injury. In particular, the uppermost layer of differentiated keratinocytes in the epidermis express caspase-8 that has a non-canonical role in regulating the wound-healing program. We previously demonstrated that the downregulation of caspase-8 is a natural phenomenon upon application of an excisional wound to the mouse skin (12). This downregulation is particularly relevant as genetically ablating caspase-8 in the epidermis is sufficient to induce a wound healing response even in the absence of any damage to the organ. In addition, the downregulation of caspase-8 in the upper, differentiated, layer of the epidermis mediates signaling networks to incite epithelial stem cell proliferation in the epidermis (12) and the hair follicle (13, 14) to fuel wound closure. We have thus used the downregulation of caspase-8 as a cellular biomarker to identify the higher order regulatory machinery that reprograms the cell to enter the wound healing process in differentiated keratinocytes, which are emerging as an important participant in the tissue repair program.

### Wound induce downregulation of caspase-8 RNA correlates with the degree of promoter methylation

Previously, we have established the importance of the downregulation of caspase-8 RNA in both physiological (wound healing (12)) as well as pathological (atopic dermatitis (15) and psoriasis (16)) scenarios. The mechanisms responsible for this downregulation, however, remains unknown. Uncovering the regulatory machinery of caspase-8 RNA also holds the promise of understanding the process by which cells transition from a state of homeostasis to repair. Moreover, it can provide potential new therapeutic targets for common inflammatory skin diseases where this regulation is perturbed.

RNA downregulation can be achieved either via blocking the synthesis and/or active degradation. In order to distinguish between these two possibilities, we determined the half-life of caspase-8 in homeostasis compared to wound conditions. In differentiated primary epidermal keratinocytes, we observed that the half-life of caspase-8 mRNA under homeostatic conditions in vitro is approximately 2 hours (Figure S1A). In an in vitro scratch wound assay with multiple scratches, the level of caspase-8 RNA is significantly reduced by 8 hours (Figure 1A). Since the reduction of caspase-8 is faster under homeostatic conditions compared to the wound healing context, merely blocking RNA synthesis can achieve the reduction of caspase-8 mRNA and initiate the downstream wound healing response. Interestingly, the reduction caspase-8 RNA is localized in cells near the front of the scratch wound in vitro (Figure 1B, Figure S1B). In situ hybridization of caspase-8 RNA demonstrates that the downregulation can clearly be visualized in the cells immediately adjacent to the leading edge of a single scratch wound as early as 4 hours post wounding. By 8 hours post-wounding, the caspase-8 RNA is downregulated in about 3-4 cell layers from the wound front. These findings are consistent with our observation in excisional wounds on the back skin of mice where the decrease of caspase-8 RNA is visible as early as 4 hours in the wound proximal region (Figure 1C and Figure S1C). Together these results suggest that simply blocking transcription post injury is sufficient to downregulate caspase-8. We hypothesized that the block in caspase-8 RNA synthesis is achieved through promoter methylation, which is consistent with previous reports documenting the same phenomenon in a variety of cancer cells through the hypermethylation of regulatory DNA sequence(17, 18). To understand whether this process in cancer cells is an aberration of the physiological healing program, we have assessed the methylation status of important regulatory sequences in the caspase-8 promoter, namely the CpG loci and SP1 binding sites (Figure S1D) (19). Analysis of methylation of SP1 sites and other CpG loci reveals a time-dependent increase of promoter methylation in a sheet of differentiated epidermal keratinocytes subjected to multiple scratch wounds (Figure 1D). This progressive increase in the methylation of the caspase-8 promoter correlates well with the kinetics of the decrease in caspase-8 RNA (Figure 1A-C). This suggest DNA methylation may play a critical role in regulating the wound-healing response.

**Figure 1:**
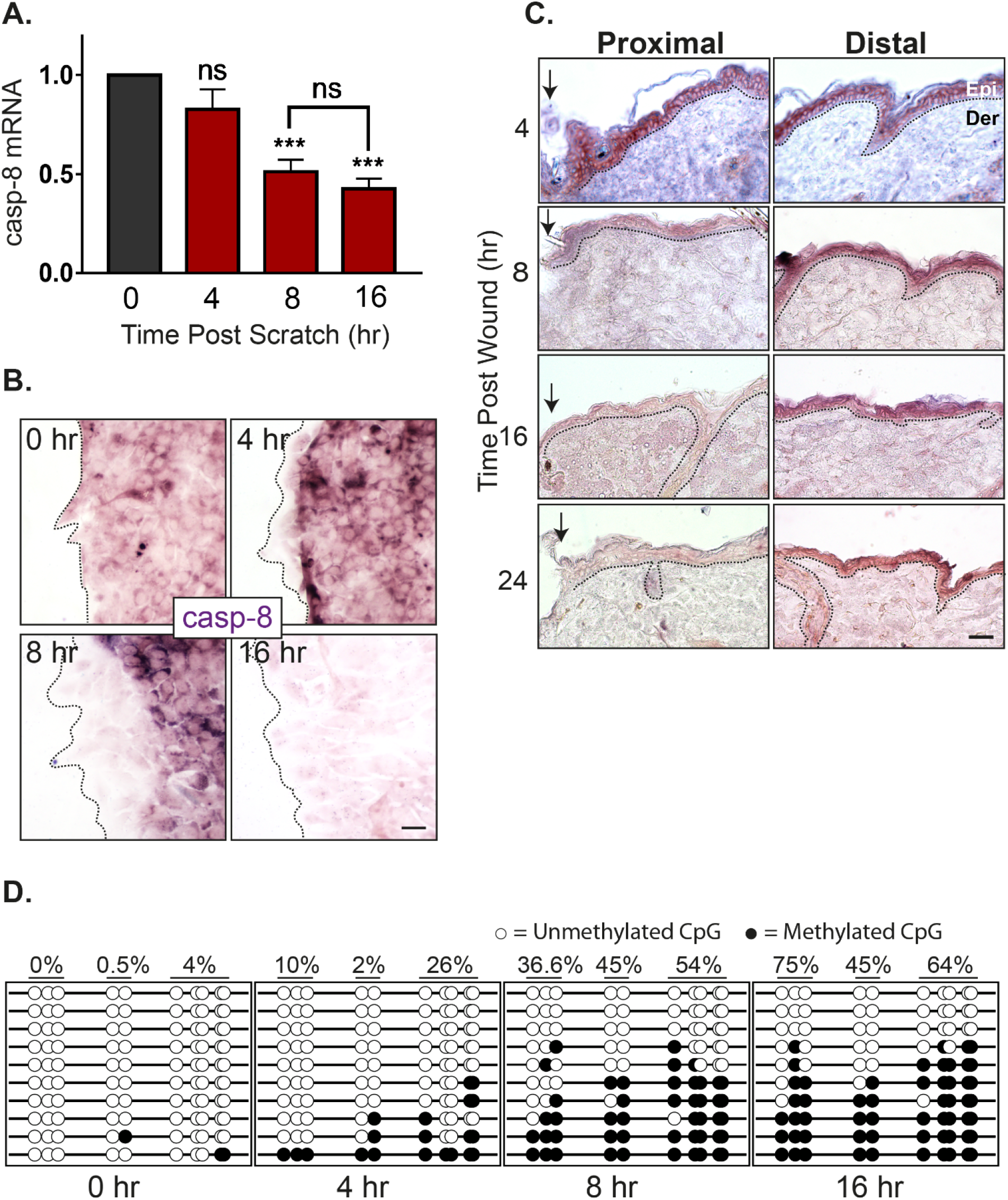
Kinetics of caspase-8 promoter methylation and expression: **A**, Levels of caspase-8 mRNA at different time-points post scratch wound (fold change) (n=4) **B**, In-vitro ISH of caspase-8 mRNA showing its levels at scratch-margins over time [scale = 10 μm] **C**, In-vivo ISH of caspase-8 mRNA showing its levels at wound proximal and distal regions over time (dotted line represents basement membrane, Epi = Epidermis, Der = Dermis) [scale = 20 μm] **D**, Bisulphite sequencing of caspase-8 promoter proximal region (265bp) shows methylation status of 10 individual CpG sites (columns) from 10 cloned PCR products (rows) at various time-points post scratch wound. Percentage value denotes the percent methylation for each group of CpG sites over time (Refer Figure S1D for the sequenced region and primer sites). (Data are shown as mean ± SEM, P values were calculated using one-way ANOVA with Dunnett’s test and two-tailed t test (A), *** P ≤ 0.001, ns = P > 0.05)

**Figure S1:**
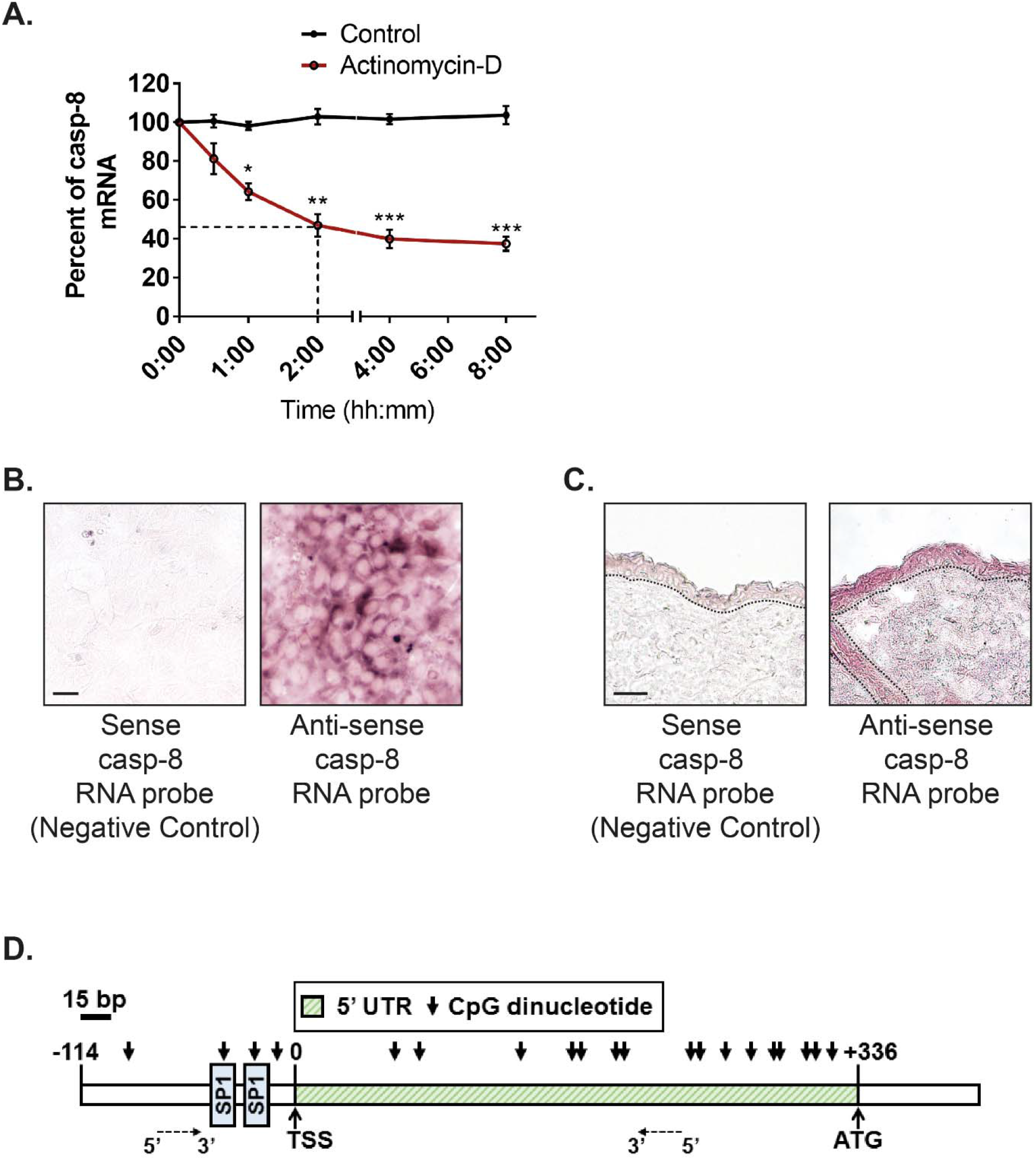
caspase-8 RNA half-life and CpG positions on its promoter proximal region. **A**, Quantification of caspase-8 mRNA to check its half-life post transcriptional block (using Actinomycin-D) **B**, In-situ hybridization with anti-sense and sense probe of caspase-8 RNA (in-vitro) [scale = 10 μm] **C**, In-situ hybridization with anti-sense and sense probe of caspase-8 RNA (in-vivo) [scale = 20 μm] **D**, Model showing positions of CpG dinucleotide and SP1 binding sites in caspase-8 promoter proximal region (Data are shown as mean ± SEM, P values were calculated using one-way ANOVA with Dunnett’s test (A), * P ≤ 0.05, ** P ≤ 0.01, *** P ≤ 0.001, ns = P > 0.05)

### Wound stimuli induce the nuclear localization of the DNA methyltransferase DNMT3a

We thus investigated the mechanism responsible for DNA methylation of the caspase-8 promoter in response to injury. The bisulfite sequencing data reveals that the methylation of the caspase-8 promoter is a de novo event in response to wounding. We therefore examined the status of the two known de novo DNA methyltransferases (DNMTs), namely DNMT3a and DNMT3b, in response to injury. Interestingly, de-novo DNMTs (DNMT3a and 3b) have also been shown to be important in regulating epidermal stem cell homeostasis (4). To investigate whether these enzymes likewise play a role in tissue repair, we examined their expression in the wounded epidermis. Consistent with a previous report, under homeostatic conditions, we found that DNMT3a mainly resides in the nucleus of the basal/proliferating (K5 positive) cells and is absent or cytoplasmic in the suprabasal/differentiated (K5 negative) keratinocytes (Figure S2A-B) (20). This localization was also recapitulated in vitro wherein we observed the cytosolic localization of DNMT3a in differentiated primary epidermal keratinocytes (Figure S2C). Interestingly, in vivo we observed that DNMT3a undergoes cytoplasmic to nuclear translocation in cells adjacent to the wound (Figure 2A). Quantification of the nuclear vs. cytoplasmic localization of DNMT3a revealed a time dependent accumulation of the enzyme in the nucleus post wounding (Figure 2B). This phenomenon was more apparent in an in vitro scratch assay, where keratinocytes adjacent to the scratch exhibited nuclear localization of DNMT3a (Figure 2C). The second known de-novo DNMT, DNMT3b, also showed cytoplasmic localization in differentiated keratinocytes (Figure S2D). However, it did not translocate to the nuclei of scratch proximal keratinocytes (Figure S2E).

**Figure 2:**
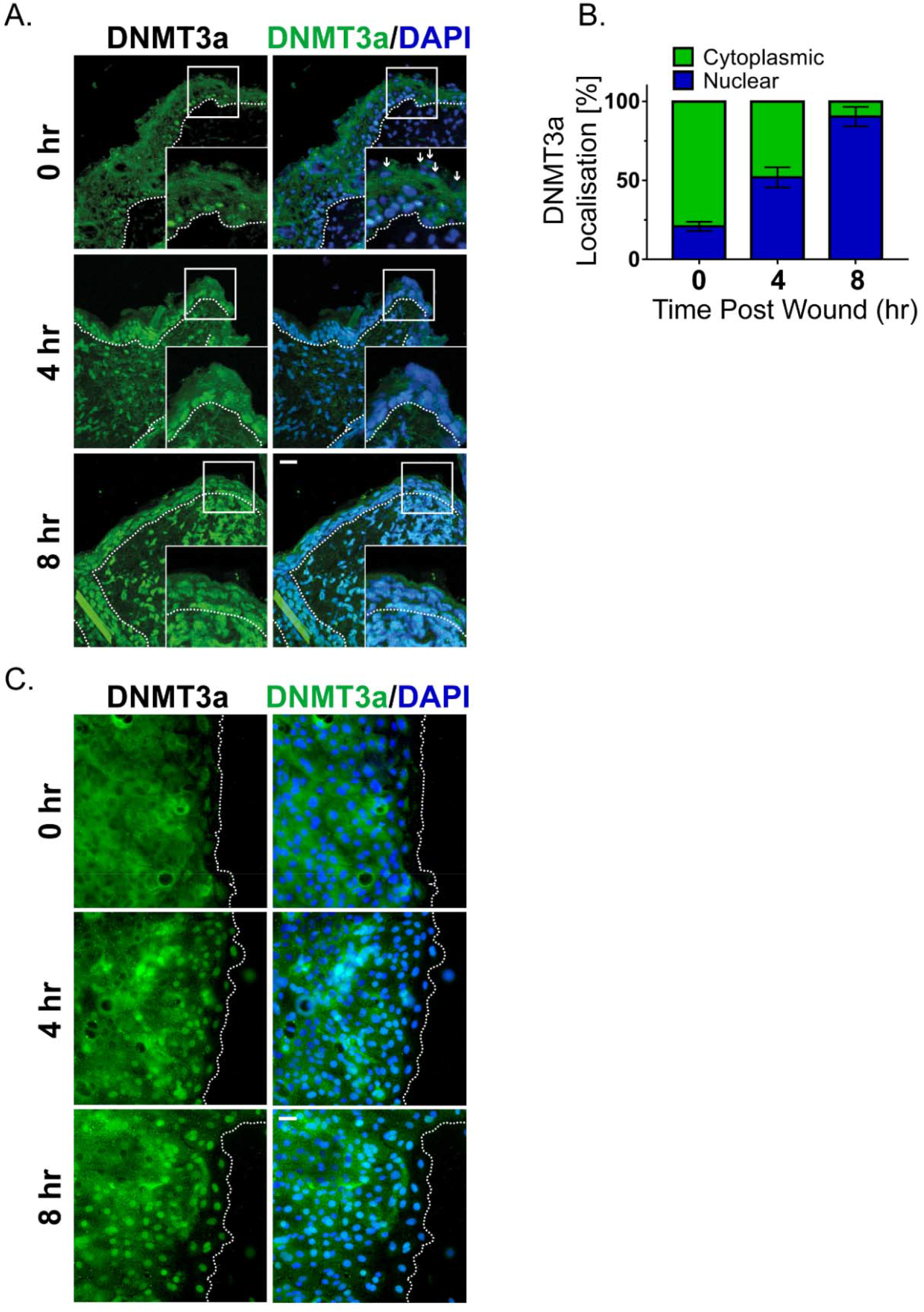
**A**, DNMT3a and DAPI staining of wound proximal (<0.5 mm) skin sections at different time interval (small white arrows showing nuclei of suprabasal keratinocytes, negative for DNMT3a staining. **B**, Quantification and kinetics of DNMT3a localization (nuclear v/s cytoplasmic) from wound proximal (≤100mm) skin sections. (it represents quantification of differentiating keratinocytes from the skin sections of three separate biological replicates) **C**, DNMT3a and DAPI staining of scratch wounded in vitro differentiated keratinocyte layer [scale = 20 μm]

**Figure S2:**
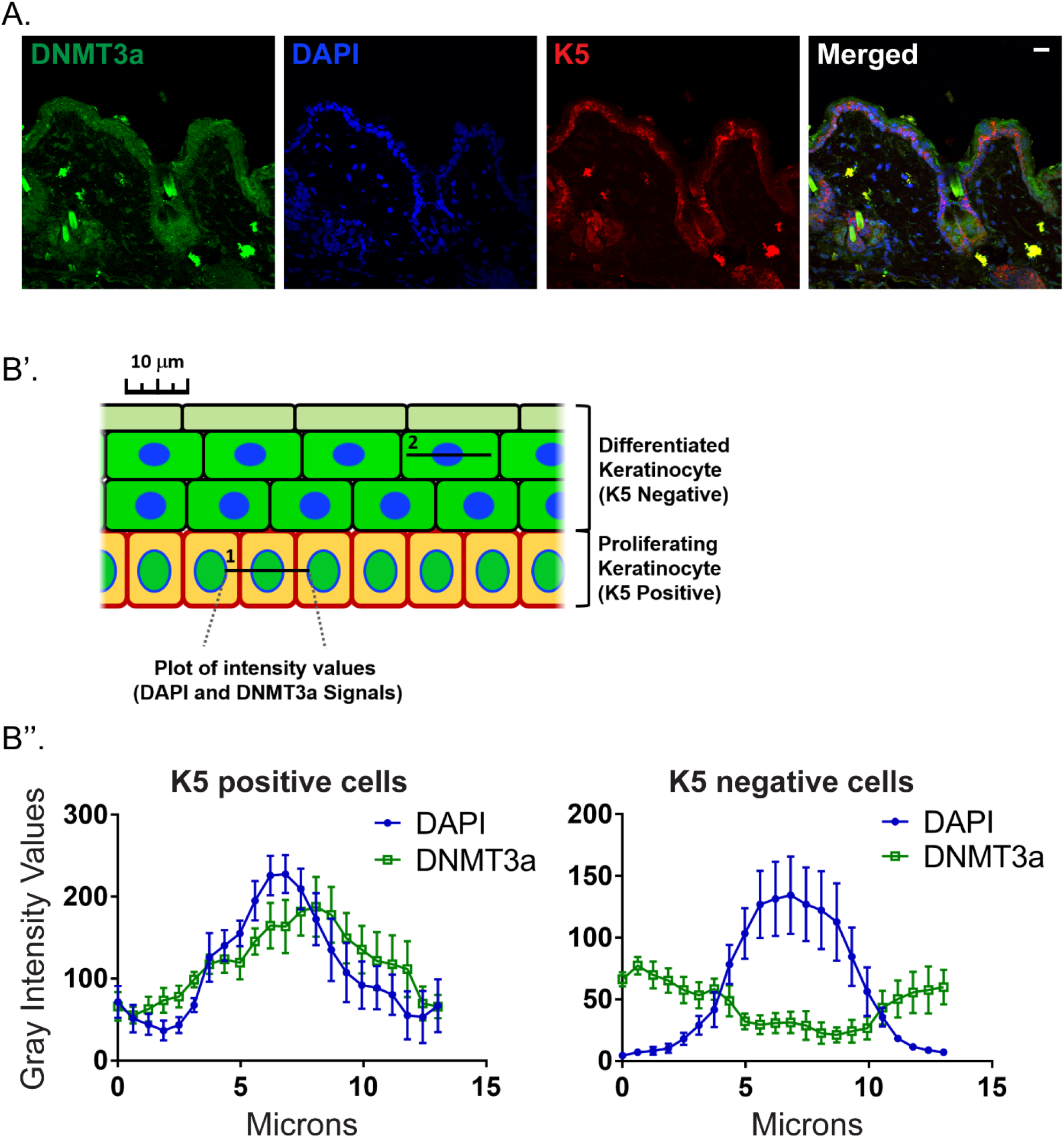
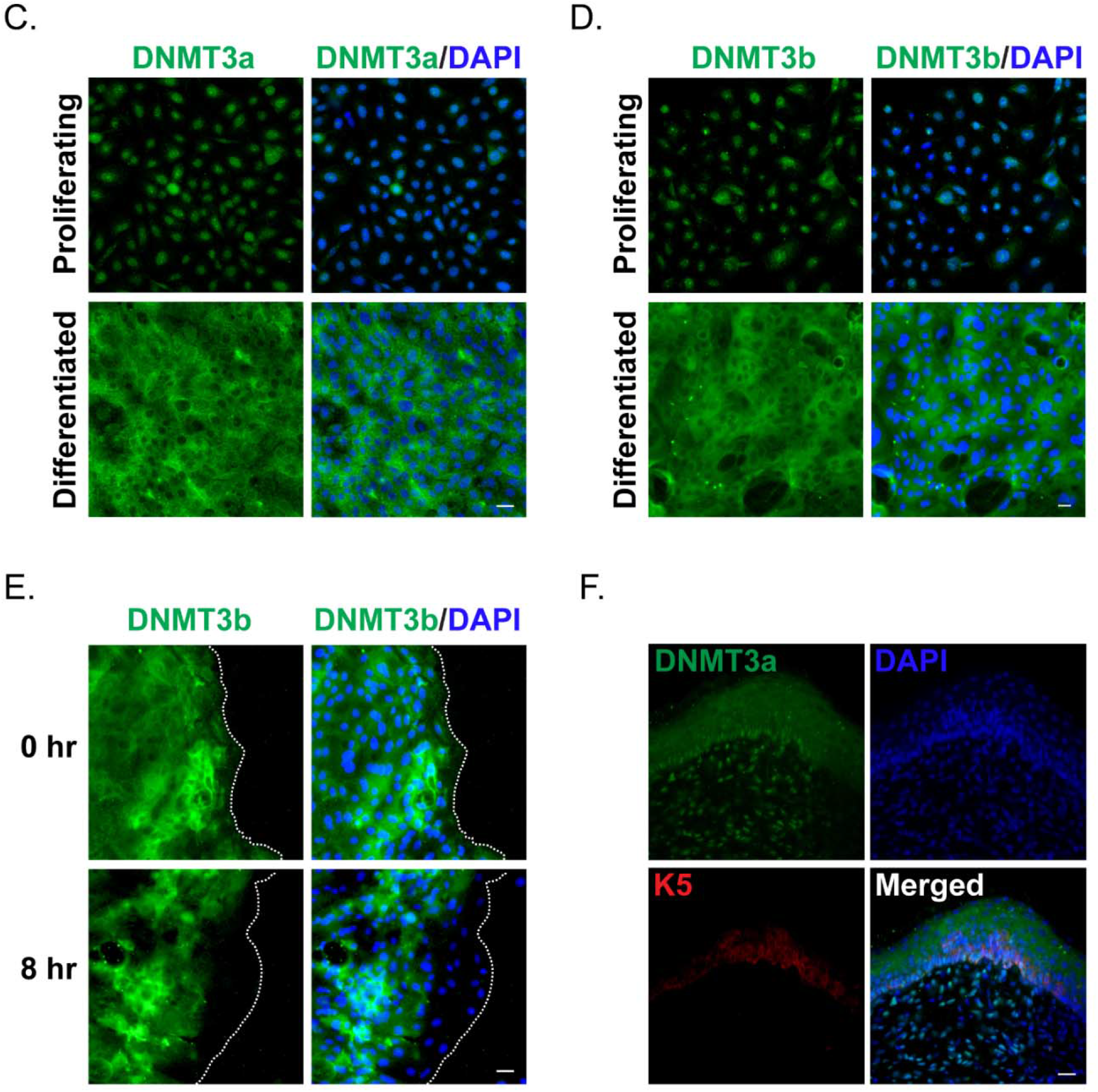
**A**, Representative image of unwounded/wound-distal skin section stained with DNMT3a, DAPI and K5. **B’**, A model showing the quantification method of DAPI and DNMT3a stain intensities over the line of interest (1, 2) from proliferating and differentiated keratinocytes, followed by **B”** the plots of intensity values (gray unit). Staining of in vitro proliferating and differentiated keratinocytes with **C**, DNMT3a/DAPI and **D**, DNMT3b/DAPI. **E**, DNMT3b/DAPI staining of scratch wounded in vitro differentiated keratinocytes. **F**, DNMT3a, DAPI and K5 staining of a completely healed mouse skin section [scale = 20 μm]

Thus, we focused on understanding the mechanistic details of DNMT3a’s role in regulating wound-healing program. The increase in DNMT3a nuclear localization was time dependent, affecting wound proximal keratinocytes first and then moves towards distal cells. At the completion of the wound-healing program we observe that DNMT3a localization is again prominent within the cytoplasms of differentiated (K5 negative) keratinocytes, while nuclear localization is restricted to cells in the basal layer of the epidermis (Figure S2F). In conclusion, we observe that the DNMT3a shows significant nuclear localisation in the wound-proximal (leading edge) cells within 4 hours of the injury and the localisation pattern further penetrates in the distal regions as time passes (Figure 2C). The nuclear localization kinetics also correlates with the pattern of caspase-8 downregulation as well as promoter methylation (Figure 1B-D).

### DNMT3a directly regulates caspase-8 expression

We further explored whether the de-novo DNA methylation of caspase-8 promoter is the result of DNMT3a’s direct binding to this region (Figure S1B). This was accomplished with the use of Chromatin Immunoprecipitation (ChIP) to assess the level of DNMT3a occupancy on the caspase-8 promoter pre- and post-scratch wound. We found that scratch wounds lead to the higher occupancy of DNMT3a on caspase-8 promoter, which is not seen in the case of DNMT3b (Figure 3A). To understand the functional relevance of DNMT activity in maintaining caspase-8 levels, we pre-treated the differentiated keratinocytes with a generic DNMT inhibitor (5-Aza-2′-deoxycytidine). We observed that the inhibitor treated cells were unable to downregulate caspase-8 mRNA in a scratch wound assay (Figure S3A). To specifically assess the role of DNMT3a, we performed shRNA mediated knockdown of DNMT3a (Figure S3B). Compared to the scrambled RNA controls, keratinocytes with reduced DNMT3a expression were unable to downregulate caspase-8 in response to scratch wound (Figure 3B). We further analysed whether failure of caspase-8 mRNA downregulation was due to the absence of promoter methylation. Indeed, scratch-wounded keratinocytes, transduced with DNMT3a shRNA, showed significantly reduced DNA methylation pattern on the caspase-8 promoter compared to scrambled RNA control (Figure 3C).

**Figure 3:**
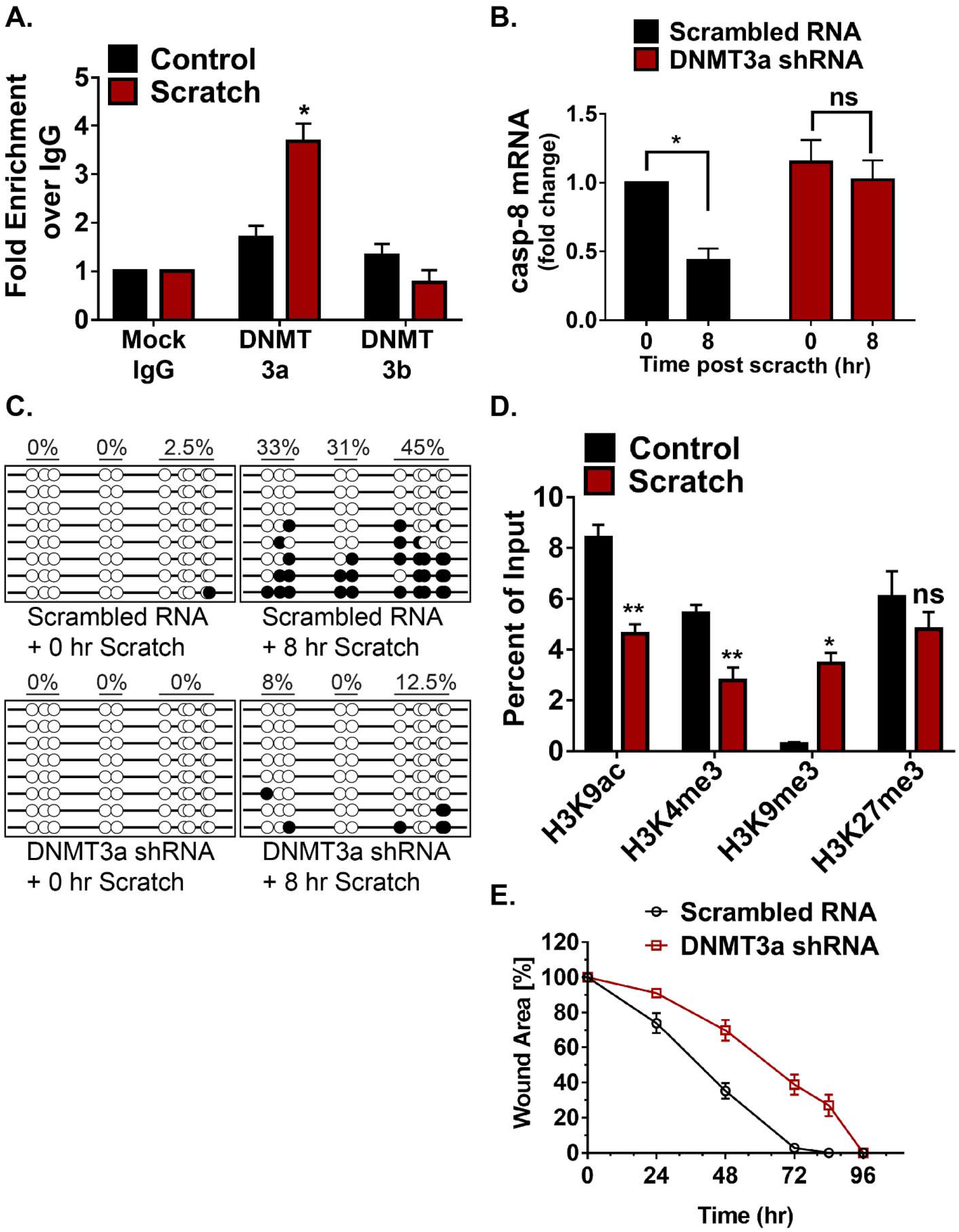
Involvement of DNMT3a and histone modification in regulating caspase-8 expression: **A**, ChIP-qPCR analysis to check DNMT3a and DNMT3b occupancy at caspase-8 promoter in control and scratch-wounded keratinocytes (n=3) **B**, qPCR analysis of caspase-8 mRNA in scratch-wounded keratinocytes, transduced with either scrambled RNA or DNMT3a shRNA (n=3) **C**, DNA methylation status of caspase-8 promoter in scratch-wounded keratinocytes, transduced with either scrambled RNA or DNMT3a shRNA **D**, ChIP-qPCR analysis of H3K9ac, H3K4me3, H3K9me3, and H3K27me3 at caspase-8 promoter in control and scratch-wounded keratinocytes (n=3) **E**, Effect of DNMT3a downregulation on in vitro wound healing assay (Data are shown as mean ± SEM, P values were calculated using two tailed t-test (A, B, D), * P ≤ 0.05, ** P ≤ 0.01, *** P ≤ 0.001, ns = P > 0.05)

**Figure S3:**
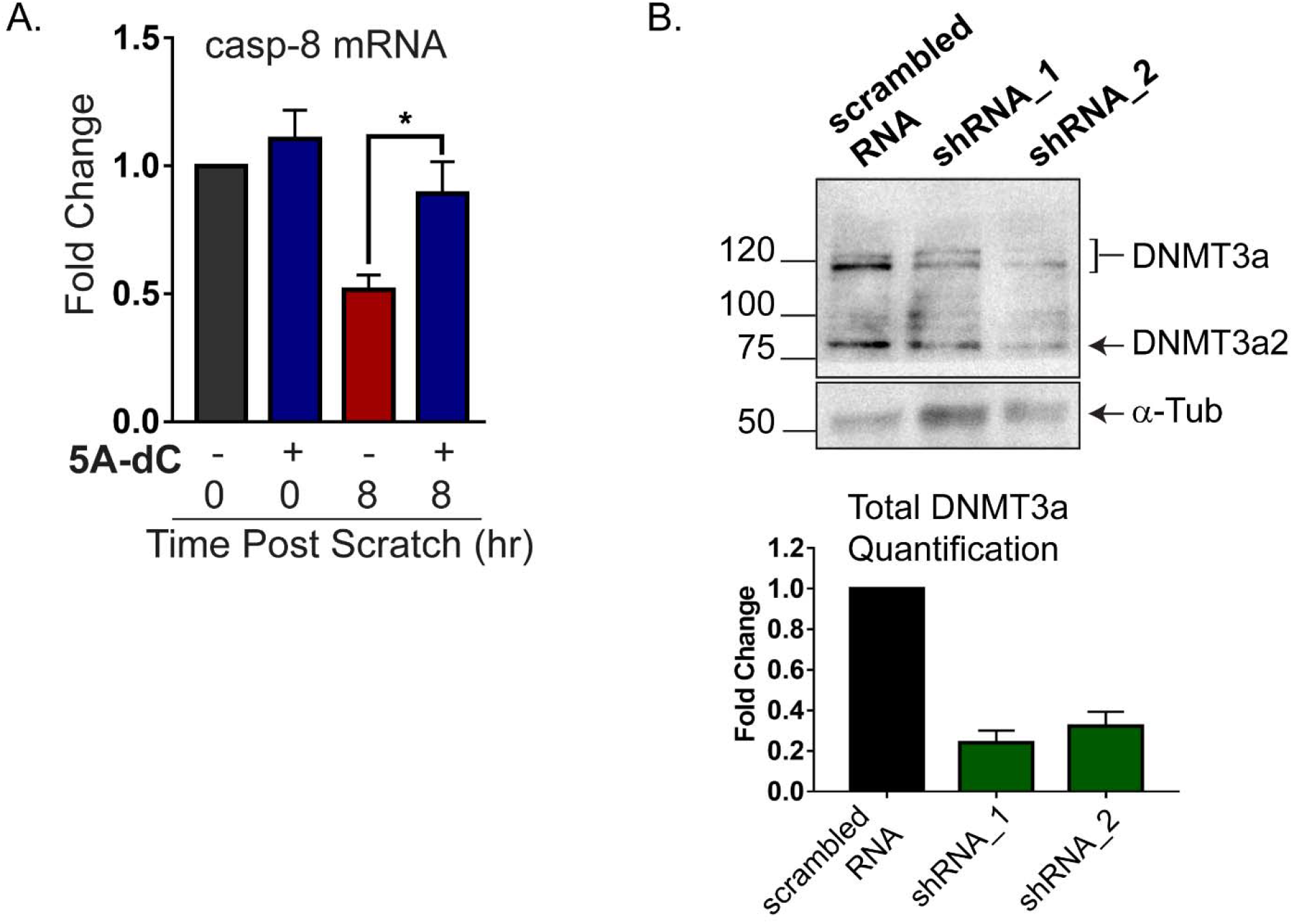
**A**, qPCR analysis of caspase-8 mRNA in scratch-wounded keratinocytes, pre-treated with 5-Aza-2′-deoxycytidine (5A-dC) or DMSO (n=3) **B**, western blot analysis from keratinocytes transduced with scrambled RNA or DNMT3a shRNA (α-Tub = alpha-tubulin) (Data are shown as mean ± SEM, P values were calculated using two tailed t-test (A), * P ≤ 0.05, ** P ≤ 0.01, *** P ≤ 0.001, ns = P > 0.05)

Promoter activities are often dependent on the associated histone modifications. These histone marks generally guide the DNA methylation at a particular genic region and vice-a-versa(21–23). DNMT3a occupancy and activity has also been shown to be influenced by the methylation status of certain lysine (K) residues on the histone 3 (H3) tail(22, 24). To investigate the core machinery required for DNMT3a mediated methylation on the caspase-8 promoter, we assessed several activation and repression histone marks in scratch wounded keratinocytes (Figure 3D). We observed that two transcriptional activation marks, H3K9ac and H3K4me3, are decreased at the caspase-8 promoter. On the other hand, the H3K9me3 mark, which is associated with transcriptional repression, was significantly increased at the caspase-8 promoter following wounding. Interestingly, another classical repressive mark, H3K27me3, did not show a significant change. It is possible that the caspase-8 proximal promoter is another example of a bivalent promoter (25) having both activation (H3K9ac and H3K4me3) and repression (H3K27me3) marks. In this scenario, then, wound-mediated repression of caspase-8 is achieved via reduction of both H3K9ac and H3K4me3 along with an increase in the H3K9me3 mark and DNMT3a occupancy. These results establish the mechanism by which DNMT3a localizes to the caspase-8 promoter. An outstanding question is whether DNMT3a is required for a proper wound healing response. To address this issue, we tested the effect of the knockdown of DNMT3a in a scratch wound assay (Figure 3E). We found that keratinocytes with decreased DNMT3a exhibited an impaired wound closure response, thereby illustrating the necessity of this methyltransferase in the proper repithelialization of an in vitro wound.

### Effect of cellular tension on DNMT3a localization and caspase-8 expression

We observed that caspase-8 downregulation and DNMT3a nuclear localization initiate at the edge of wound site (Figure 1, 2). Given that these are early responses to injury, understanding the mechanistic basis of this phenomena can provide insights into the broader process of cellular wound sensing. The keratinocytes in the epithelial sheet are strongly connected to each other and an event of injury will result in the sudden relaxation in that tension, particularly in the cells at the boundary of the wound. Interestingly the expanding number of cells exhibiting the downregulation of caspase-8 RNA in the scratch wound assay over time (Figure 1A) closely parallels the changes in traction force previously reported for the collective cell migration of an epithelial sheet following a scratch wound (26). We therefore investigated whether release of tension, caused by the severing of the epithelial sheet, can impact DNMT3a subcellular localization and subsequently caspase-8 expression. As shown in Figure S4a, modulation in cellular tension can be achieved via targeting the components of the adherens junction, which are known to play a role in generating and maintaining cellular tension (27, 28).

We observed that tension release by disrupting calcium-dependent E-cadherin junctions via EGTA treatment resulted in the nuclear localization of DNMT3a (Figure 4A). Similarly, releasing cellular tension endowed by Non Muscle Myosin II (NM-II) with the pharmacological inhibitor of NMII, blebbistatin, induced the DNMT3a’s nuclear translocation from the cytosol (Figure 4A). Furthermore, we examined the effect of blocking release of cellular tension in a scratch wounded sheet of epidermal keratinocytes. The release of tension was blocked by pre-treating keratinocytes with calyculin-A, which inhibits myosin light chain phosphotase, thereby maintaining the active state of NMII (29). The treatment of keratinocytes with calyculin-A prior to scratch wounding blocked the nuclear translocation of DNMT3a that was observed in cells treated with vehicle control (Figure 4B).

**Figure 4.**
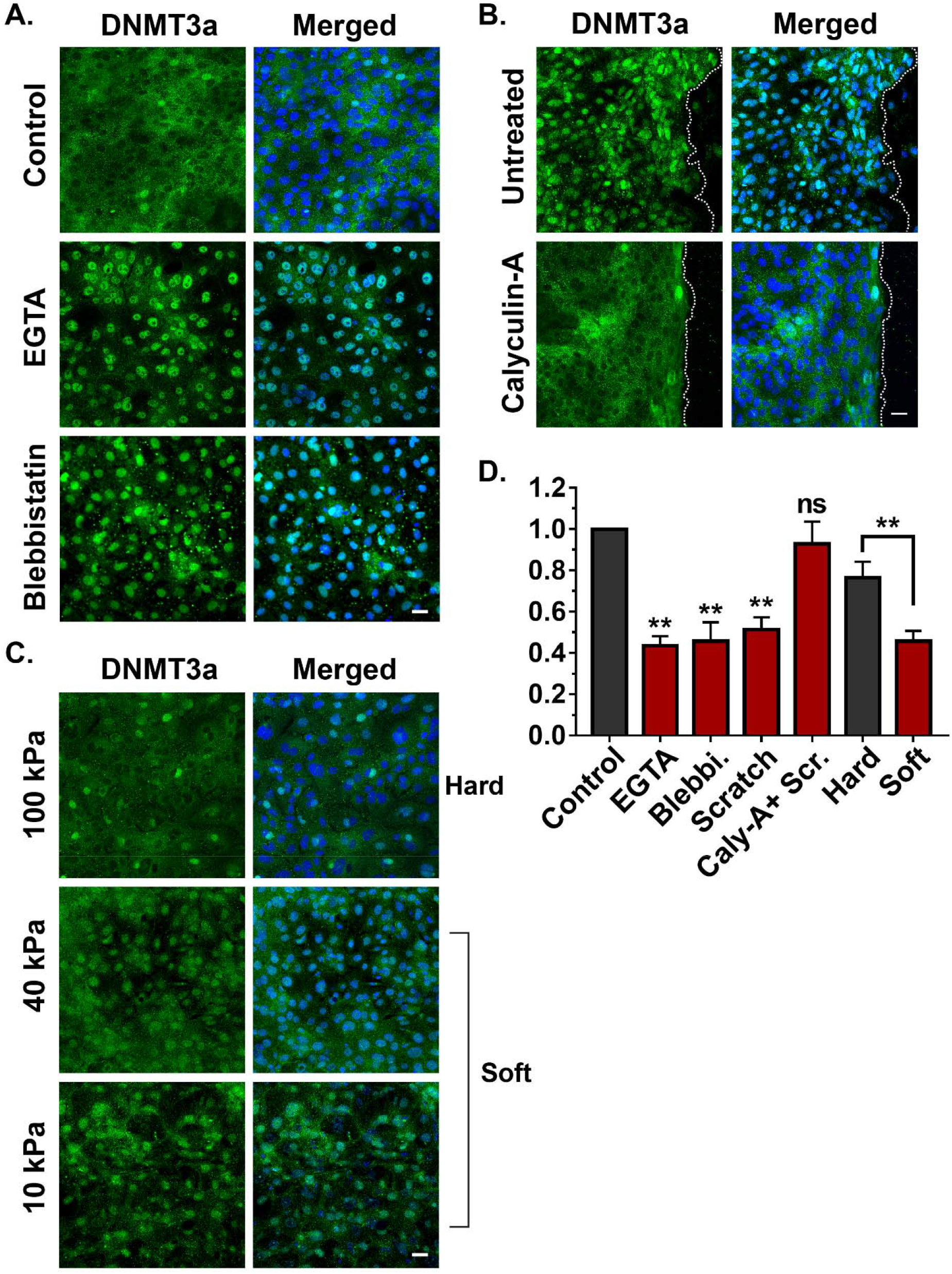
Effect of cellular tension on DNMT3a localization and caspase-8 expression: **A,** Effect of EGTA and Blebbistatin on the localisation of DNMT3a **B**, Effect of scratch wound DNMT3a localisation in presence and absence of calycuilin-A **C**, Effect of various matrix stiffness on the localisation of DNMT3a **D**, Fold change in the levels of caspase-8 mRNA as a result of varios pharmacological and mechanical approaches of tension modulation (n=3), [scale bar = 20 μm] (Data are shown as mean ± SEM, P values were calculated using two tailed t-test (D), * P ≤ 0.05, ** P ≤ 0.01, *** P ≤ 0.001, ns = P > 0.05)

**Figure S4:**
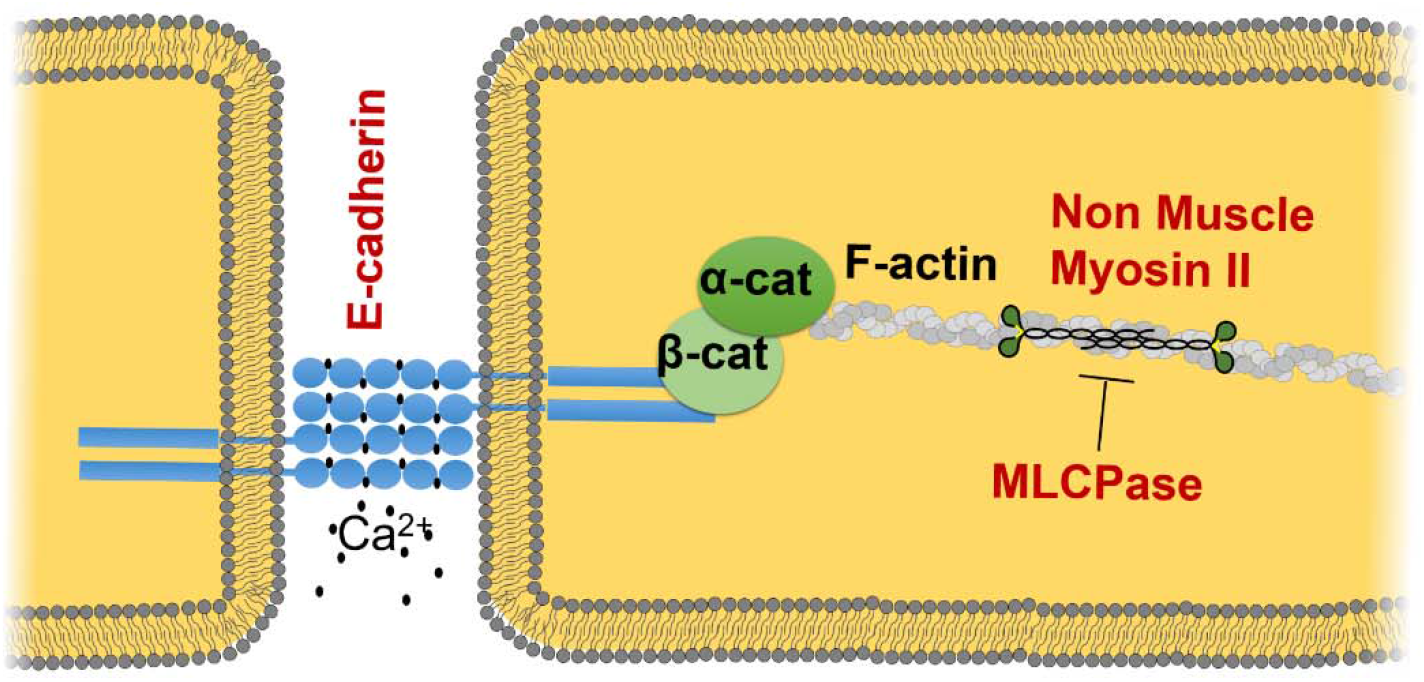
Model showing various potential protein molecules (red labels) involved in generating and/or sensing the cellular tension

In addition to a pharmacological approach, we also modulated cellular tension by altering the substrate stiffness on which the keratinocytes were growing. This was accomplished by utilizing polyacrylamide gels of various stiffness, which would alter cellular tension. We observed that differentiated keratinocytes seeded on “soft” matrices ranging from 10 kPa to 40 kPa mostly harboured DNMT3a in the nuclei (Figure 4C). However, cells grown on a “stiffer” matrix (100kPa) predominantly showed a cytoplasmic localization of DNMT3a.

We then evaluated whether DNMT3a’s dynamic localization in response to pharmacological and mechanical alterations in cellular tension has any transcriptional consequences. We observed that in all the scenarios where DNMT3a nuclear localization was favored (scratch wounds, EGTA/blebbistatin treatment, soft substrates), caspase-8 RNA was downregulated compared to their respective controls (Figure 4D). On the otherhand, inhibition of DNMT3a’s nuclear localization (via calyculin-A, or a stiff substrate) resulted in the failure of caspase-8 downregulation in spite of a scratch wound. These results suggest that keratinocytes organized within an epithelial sheet can translate changes in tensile forces into cellular reprograming via epigenetic means.

### DNA methylation could be a global regulator of gene expression to initiate wound-healing program

We further assessed whether the downregulation of caspase-8 is a paradigm for the global downregulation of genes to achieve a cell state transition from homeostasis to wound healing. Surprisingly, the transcriptome profile of scratch wounded differentiated keratinocytes has not been reported even though these layers are the first to encounter damage in vivo. Thus we performed RNA sequencing of wounded v/s unwounded primary mouse keratinocyte that were differentiated via the calcium switch protocol (Figure 5A). The analysis of the transcriptome data revealed that the number of downregulated genes outnumbered the upregulated genes post injury. We verified the sequencing data by specifically analysing genes via qPCR that have already been implicated in wound healing or epidermal development (Figure 5B and C). Interestingly, there was an inverse correlation with the RNA expression and the degree of methylation for many of the genes we interrogated. This suggest that DNA methylation could be a global regulator for a set of wound response genes (in addition to caspase-8), needed for the wound healing program.

**Figure 5:**
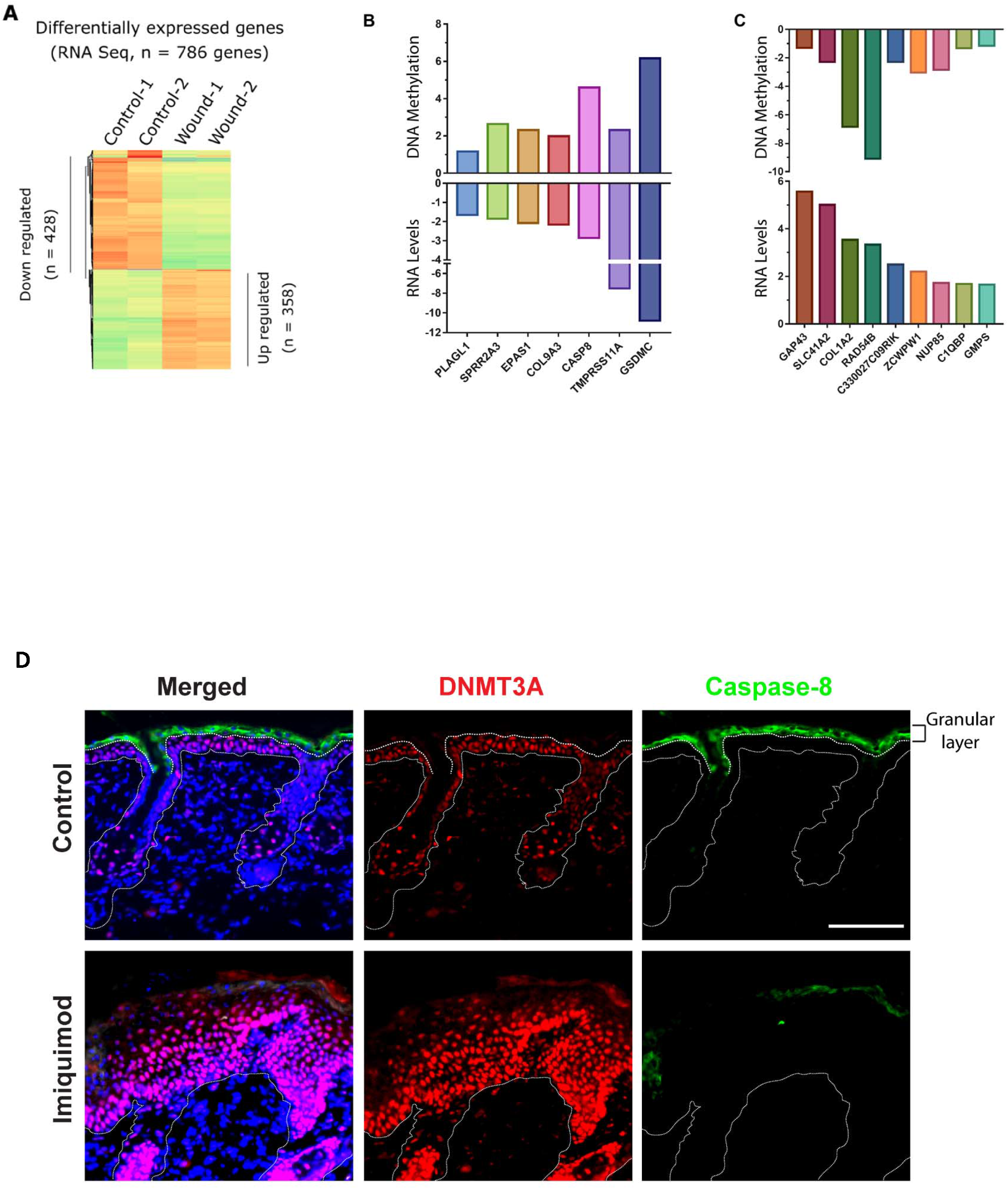
**A**, Heat map of differentially regulated genes in Control and Scratch wounded keratinocytes. **B**, Transcriptionally downregulated genes and their associated DNA methylation levels **C**, Transcriptionally upregulated genes and their associated DNA methylation levels. (MeDIP-qPCR, y-axis = fold change compared to control) **D**, DNMT3a and caspase-8 staining of Control and Psoriatic mouse skin (induced through imiquimod treatment). [scale bar = 100 μm]

**Figure S5:**
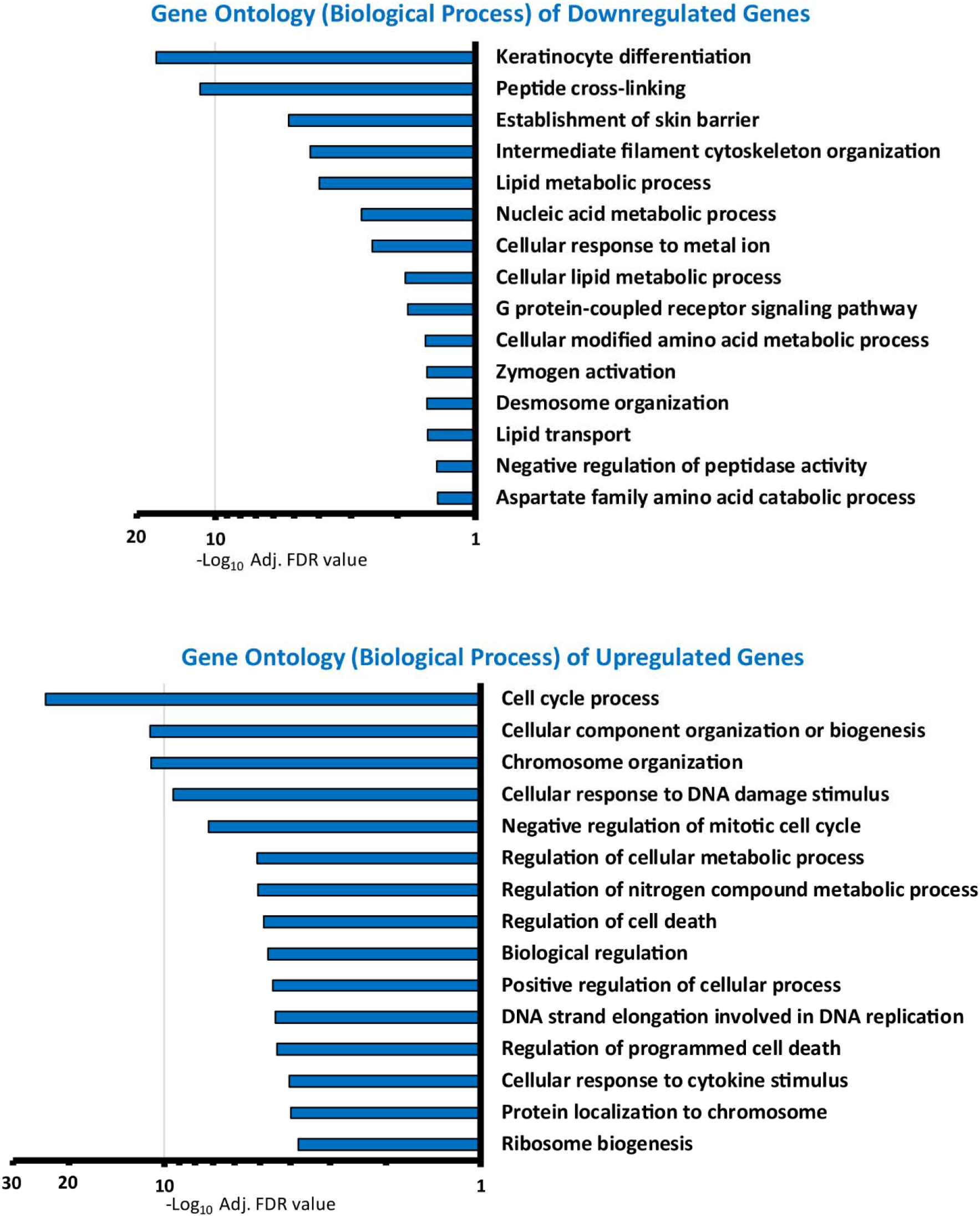
Gene ontology of Upregulated and Downregulated genes (Biological Processes). Processes are listed as –Log_10_ of adjusted FDR values. Top 15 relevant biological processes are chosen for generating the graphs.

Analysis of transcriptome data has revealed many such group of genes and their biological processes. (Figure S5). Of particular interest were the downregulation of genes involved in the differentiation of keratinocytes. The Ghazizadeh lab has reported evidence of dedifferentiation of suprabasal keratinocytes as a mode of aiding cutaneous regeneration and repair. Interestingly, the regeneration of skin epithelia by differentiated epidermal cultures was found to be facilitated by the capacity of these cells to proliferate (30). The transcriptome profile of scratch wounded differentiated keratinocytes reveals an upregulation of cell cycle associated genes that is consistent with this report. Consequently, the convergence of mechanical and epigenetic cues appears to play an important role in the plasticity of differentiated epidermal keratinocytes in cutaneous repair and regeneration. The processes that occur during the wound healing phases of inflammation, proliferation, and tissue remodeling are often reproduced in a deregulated manner in many pathologies leading to the notion of diseases with a “wound signature”. Prominent among these is the view of cancer as an overhealing wound (31). As we noted earlier, there is a body of literature demonstrating that the downregulation of caspase-8 in cancer cells is accompanied with the methylation of its promoter region (32–35). In addition, we have previously demonstrated that inflammatory human skin diseases such as atopic dermatitis (15) and psoriasis (16) likewise exhibit a loss of epidermal caspase-8. To probe a possible link between caspase-8 downregulation and methyltransferase expression, we utilized the imiquimod-induced model of psoriasis in mice. In the psoriatic skin of mice, we observed robust nuclear localisation of DNMT3a in all the epidermal layers, whereas in the control animals, nuclear DNMT3a was primarily localized in the basal keratinocytes (Figure 5D). Altogether, this suggests that the epigenetic regulation governing the cell state transition in wound healing is usurped in many diseases ranging from inflammatory skin diseases to carcinomas.

## DISCUSSION

The wound-healing literature involving epidermal keratinocytes have elegantly described many signaling pathways and gene expression profiles in the proliferating cells of the basal layer (36, 37). In contrast, differentiated epidermal cells, such as the suprabasal keratinocytes near the outer surface of the skin, have largely been overlooked for their potential role during wound-healing. Interestingly, our previous work demonstrates that the uppermost layer of differentiated keratinocytes, namely the granular layer, expresses caspase-8 that has a non-canonical role in regulating the wound-healing program (12). It was found that the downregulation of caspase-8 is both necessary and sufficient to induce a wound healing response in the absence of any tissue damage. In addition, the chronic downregulation of caspase-8 underlies inflammatory skin diseases such as atopic dermatitis (15) and psoriasis (38). These findings have made the decrease in caspase-8 expression a useful wound-healing biomarker and led us to inquire about the mechanism of caspase-8 regulation in skin keratinocytes.

Clues about the regulation of caspase-8 is reported in the context of cancer. Similar to the wound-healing process, it is generally downregulated in various cancers (18, 39, 40). It is possible that the cancers, known as “over healing wound”, usurp physiological pathway of wound-healing for its own propagation (31). Here we show that wound-sensing leads to the acute increase in the caspase-8 promoter methylation as a potential mechanism of gene silencing. This parallels with the findings on caspase-8 downregulation in hepatocellular carcinoma, where methylation status of SP1 sites and nearby CpG dinucleotides in the promoter region were proposed to be a major regulator of caspase-8 expression (19). In fact, it has been observed that caspase-8 and several other genes are known to be downregulated in various cancers via DNA methyltransferase (DNMT) activity (18, 39, 40). The overexpression of DNMT3a has also been shown to be associated with several cancers (41, 42)The process of de-novo DNA methylation during an acute physiological response such as wound healing is a rarely described phenomenon. Mammalian cells are known to have only two de-novo DNA methyltransferases, DNMT3a and DNMT3b. Both have been widely studied for their role in physiological processes like embryogenesis (43) and hematopoiesis (44), as well as pathological conditions such as cancer (45, 46). In particular, , it has been shown that DNMT3a and 3b are required as regulators of enhancer activity and RNA production of genes necessary for epidermal stem cell homeostasis (4). In disease context, DNMT3a has been described to be overexpressed or mutated in various carcinomas (47, 48) and correlates with the downregulation of caspase-8 in these same scenarios. Here, we found that upon injury to the skin or differentiated epidermal sheets, the suprabasal cells near wound edge showed a nuclear localization of DNMT3a, but not DNMT3b. We have captured that DNMT3a indeed occupies the caspase-8 promoter and plays an important role in its downregulation post injury. In parallel to the DNA methylation, the literature also describes changes in histone modifications responsible for the ON/OFF state of a particular gene. The histone modifications and their modifiers have been studied in depth to understand how the expression of various epidermal differentiation genes are regulated (49). In general, H3K9ac, H3K4me3 are considered as gene activation marks and H3K9me3 and H3K27me3 are known as repression mark. It is also observed that certain methylation state of H3K36 dictates the DNMT3a’s recruitment to a particular DNA segment on the chromosome (50, 51). In our efforts to understand the histone modifications during wound-healing, we observed a reduction in H3K9ac and H3K4me3 levels, along with an increase in the H3K9me3 mark at the caspase-8 promoter. These histone modifications are known to be regulated via various other epigenetic players such as Polycomb repressive complexes (PRC 1/2), JMJD, Setd8, and HDACs during epidermal development (49).

How these epigenetic players are regulated is another important question in the field. While there are many chemical cues, adhesion signals, and transcription factors described to regulate the wound-healing process, emerging evidence links mechanical forces to epigenetic and transcriptional responses (52, 53). Even during the development of epidermal tissue, tension generating molecular players like non-muscle myosin IIA (NMIIA), along with emerin (Emd) and PRC2 regulate the differentiation process of epidermal stem cells. The strain on epidermal cells reduces Emd levels from the inner nuclear membrane, which then leads to the loss of the histone mark H3K9me2,3. This is followed by Polycomb repressive complex 2 (PRC2) mediated increase of H3K27me3 occupancy at several heterochromatic regions and thereby gene silencing (54). Along the same line, recently Nava et al. has described how short and long term mechanical stress on a cell can result in changes in stiffness of the nuclear membrane, loss of H3K9me3 marks at the heterochromatin and overall chromatin and cytoskeletal reorganization (55). These are some of the key discoveries suggesting external mechanical forces drive changes in heterochromatin organization, gene expression changes, and cytoskeletal reorganization in a way that mechanical energy gets redistributed and DNA damage can be avoided. In this context, our results demonstrate that the release in the mechanical tension, either by physical or chemical treatments, results in the DNMT3a’s nuclear localization and downregulation of caspase-8. This observation is consistent with the concept of mechano-sensitive histone modifications, which could lay a foundation for the occupancy of DNMT3a. In a wider context of cellular reprogramming during the wound response, mechanotransduction seem to have a large impact on the transcriptome of the cell via the concomitant initiation of several epigenetic pathways. Future studies in this area will include elucidation of the connection between the release of mechanical tension and their sensing by these epigenetic machineries. For example, DNMT3a has been shown to have multiple binding partners (DNMT3L, SUMO-1, Cbx4, Ubc9, RP58, HDAC1) for their nuclear shuttling as well as chromosomal occupancy, some of which can potentially function as a primary signal sensor to guide the localization of DNMT3a (56, 57). Moreover, in different cell types, changes in mechanical tension have been documented to directly induce the nuclear translocation of important transcription factors. A notable example of which is the YAP/TAZ complex, which has proliferation stimulating gene targets (58).

The described model of mechanosensitive epigenetic players would obviously be regulating a larger gene regulatory network, in addition to caspase-8. Interestingly the transcriptome literature on wound-healing has utilized proliferating keratinocytes, leaving the transcriptome profile of differentiated keratinocytes unknown despite the fact that it constitutes about 2/3 of the epidermis. Our research fills an important gap by providing a transcriptome profile of in vitro wounded differentiated keratinocytes. The results give us a unique insight in the regulation of various unexplored wound-response genes. On a particular note, we observe a strong downregulation of multiple epidermal differentiation genes in response to injury. From the current transcriptome and literature survey it is evident that various keratinocyte differentiation markers (such as involucrin, keratins K1/K10, and filaggrin) are downregulated along with cell adhesion molecules (involved in tight junction, adherens junctions, and desmosomes). This is consistent with a report from S. Ghazizadeh’s lab that de-differentiation of suprabasal keratinocytes is a contributing factor in the wound healing response (30). Our data suggests that the release of mechanical tension in differentiated keratinocytes is one component in this process by inducing a “partial de-differentiation” and perhaps additional soluble signaling cues are required to achieve complete dedifferentiation.

## Materials and Method

### Cell culture and scratch wound assay

The isolation of primary keratinocytes from neonatal mice was performed as described in (59). Briefly, mice pups were sacrificed and the skin was removed. The skin was kept in dispase at 4° C overnight (or 37°C for 1 hour) to separate epidermis. The epidermis was then digested with trypsin to isolate keratinocytes. These cells were filter with 70-micron mesh and cultured further as described in (Novak et al. 2009) (60). The keratinocytes were cultured in lab with feeder cells (3T3J2) for 10 passage. Then feeder independent keratinocytes were taken and tested for their differentiation potential via calcium switch protocol (61). Various differentiation markers were checked via qPCR. The batch of cells showing proper differentiation and morphology were then selected for further experiments.

Proliferating keratinocytes were maintained in low Ca^2+^ E-media (0.05mM). For differentiation they were allowed to reach 100% confluence and then introduced with high Ca^2+^ (1.2mM) E-media for 48 hrs. Once they differentiated and appear as sheet-like morphology, scratch-wounds were made (with the help of a 1ml tip) at multiple sites in each culture plate. To keep the constancy between experiments, the distance between the consecutive scratch was kept approximately 0.5 mm. The scratch wounds were followed by a 1X PBS wash, and fresh high Ca^2+^ (1.2mM) E-media were added to each plate. As described in the figure legends, the cells were harvested at several time points using TRIzol reagent for RNA isolation or using lysis buffer for DNA isolation.

### Mice

C57Bl6/J animals were originally purchased from Jackson Laboratory (Stock No. 000664) and were bred for > 10 generations in the NCBS vivarium facility. 8-week-old mice were anesthetised and 5 mm or 8 mm punch biopsies were used to make full-thickness excisional wounds.

### Tissue Section and Staining

Wounded regions were embedded in OCT, frozen on dry ice, and stored in -80° freezer for further sectioning and antibody staining. 10-15 micron section were taken, stained with primary antibody at 4° C overnight, and then with secondary antibody at RT for 20-30 minutes. Antibodies used in this study are as following; Caspase-8 (Enzo #ALX-804-447-C100), DNMT3a (abcam #ab2850, SC #365769), K5 (lab generated). Sections were imaged using IX73 Olympus microscope.

### In-situ hybridisation

DIG labelled 5’ mouse caspase-8 cRNA probe was synthesized as per the manufactures instructions (Roche dig labelling kit – # 11175025910). In situ hybridisation was performed as described earlier (62). Briefly, the paraffin tissue sections were deparaffinised by treatment by xylene and ethanol gradient, or the 4%PFA fixed cells were permeabilized using 0.2% TritonX-100 for 10 minutes at room temperature. 5ng DIG labelled cRNA probes per 100uL hybridisation buffer was applied on the sections overnight at 63^0^C. Same concentration of DIG labelled mRNA with the complimentary sequence to cRNA was used as a negative control. Washing was done at 65^0^C. The Anti-DIG antibody (Roche # 11093274910 Roche) was applied overnight as per manufacturer’s instructions. Sections were developed for 30 minutes at 37^0^C using BCIP/NBT solution (Sigma # B6404). Reaction was stopped using de-ionised water once the purple colour was developed. Sections were mounted using MOWIOL solution and imaged using bright-field microscope.

### Hydrogel of varying stiffness

The polyacrylamide based hydrogels were prepared as describe in (63) and (64). They were coated with collagen and seeded with enough cells to make it 80-100% confluent, and were allowed to settle for 24-48 hr before initiating the keratinocyte differentiation.

### Chromatin Immunoprecipitation (ChIP)

ChIP of histone modification was performed as described previously (65) with some modifications. In brief, harvested keratinocytes (unscratched and scratched) were cross-linked with 1% formaldehyde. Cells were lysed in buffer N containing DTT, PMSF, and 0.3% NP-40. After isolation of nuclei, chromatin fractionation was done using 0.4 U of MNase (N5386, Sigma) at 37°C for 10 min. Reaction was stopped using MNase stop buffer without proteinase K. Simultaneously, antibodies against H3K27me3, H3K9me3, H3K4me3, H3K9ac and Rabbit IgG were kept for binding with Dynabeads for 2 hr at RT. After equilibration of beads, chromatin was added for pre-clearing. To antibody bound beads, pre-cleared chromatin was added and kept for IP at 4°C overnight. Next day, beads were washed and eluted at 65°C for 5 min. Eluted product was subjected to reverse cross-linking along with input samples, first with RNase A at 65°C overnight and then with Proteinase K at 42°C for 2 h. After reverse cross-linking, DNA purification was performed using phenol:chloroform extraction method. Antibodies used for this protocol are listed here:

| *Antibody* | *Cat #* | *Company* |
| --- | --- | --- |
| H3K27me3 | 07-449 | Millipore |
| H3K9me3 | ab8898 | abcam |
| H3K9ac | ab4441 | abcam |
| H3K4me3 | ab8580 | abcam |

### Bisulphite reaction, sequencing, and analysis

Genomic DNA was isolated by salting out method as described elsewhere (66), then treated with RNase for 1 hour at 37° C. Further, ∼ 20 microgram DNA was taken in 200 microliter volume and purified with phenol:chloroform extraction method. The purified DNA was checked for its integrity via running on the agarose gel. The DNA sample having good integrity and free of RNA were taken for bisulphite conversion as per manufacturer’s protocol (Zymo #D5005). The converted DNA was then amplified using bisulphite conversion specific primers, the amplified product was assessed on the agarose gel, ligated with TOPO-TA vector.

and then sent for Sanger’s sequencing. caspase-8 promoter - for bisulphite sequencing (GAATAAGGAAGTGTTTTTTAG, AAAACTATACTCACTTCCTATTC). The sequenced file (FASTA) was uploaded to http://quma.cdb.riken.jp/ for CpG methylation analysis.

### Lentivirus shRNA constructs and transduction

Plasmids expressing shRNAs were obtained from TransOmics (DNMT3a # TLMSU1400). To produce viruses, HEK293T cells were transfected with psPAX2, pMD2.G, and either non targeting random RNA sequence vector or shRNA- containing plasmids, using Lipofectamine® LTX & PLUS transfection reagents according to the manufacturer’s protocol. Following a 48-72 hr transfection, the virus particle-containing media was collected, concentrated with filters, and added to the differentiated cells for 24 hr. Expression of DNMT3a was measured two to three days after viral infection. Silencing efficiency was confirmed by immunoblotting.

### Quantitative Real Time PCR

RNA was isolated from human keratinocytes (proliferating or differentiated) using the RNAiso Plus (Takara). 1 μg of RNA was used to prepare cDNA using the PrimeScript kit (Takara). cDNA equivalent to 100 ng of RNA was used for setting up the qPCR reaction using the SYBR green 2x master mix. All reactions were performed in technical triplicates using the CFX384 Touch Real time PCR detection system (BioRad). Primers used in this study are listed here: caspase-8 mRNA (TCTGCTGGGAATGGCTACGGTGAA, GTGTGAAGGTGGGCTGTGGCATCT), caspase-8 promoter (GGGAATAAGGAAGTGTCCTCCA, CCCAGAACTGTACTCACTTCCTG), beta Actin (GGGCTATGCTCTCCCTCAC, GATGTCACGCACGATTTCC)

### RNA Sequencing and data analysis

The scratch wounded cells and controls were collected after 8 hours in TRIzol reagent and RNA was isolated using standard TRIzol based RNA isolation method. The library preparation and NGS RNA sequencing steps were outsourced to a commercial facility (Genotypic). Once the raw sequencing reads were received, sequencing data analysis was performed using the following analysis pipeline. Briefly, raw sequencing data was QC checked with the “FASTQC” tool (Babraham Bioinformatics). Adapter contamination and bad quality reads were trimmed using “Trimmomatic” tool (67). The good quality reads were then mapped to mm10 (mouse) reference genome using “HISAT2” (68). The resulting “SAM” outputs were converted to “BAM” output and sorted. The “HTSeq-Count” tool was used to generate expression matrix from all four samples. Then, differential expression was analysed with the help of DESeq2 R package.

### Gene Ontology Enrichment Analysis

To explore enrichment of Gene Ontology among the significantly down regulated (n = 428) and up regulated genes (n = 358) we have used http://geneontology.org/ resources which runs “PANTHER” for the enrichment analysis (69).

Additional details of the NGS RNA seq samples are given here:

| Sample Names | Raw Sequencing read counts | Good quality read counts (Used for mapping to mm10 reference genome) |
| --- | --- | --- |
| Control-1 | 33693828 | 30441624 |
| Control-2 | 34605485 | 30973898 |
| Scratch-wound-1 | 32255293 | 27735961 |
| Scratch-wound-2 | 31056628 | 27761854 |

## ACKNOWLEDGEMENTS

The authors would like to thank Prof. Apurva Sarin, Prof. Tapas Kundu, Prof. Sudhir Krishna and Jamora Lab members for their critical review of the work and insightful discussions. This work was supported by grants to C.J. from the Department of Biotechnology of the Government of India (BT/PR8738/AGR/36/770/2013 and DBT/PR32539/BRB/ 10/1814/2019); and inStem Core funds. TB was supported by PhD scholarship from Council of Scientific & Industrial Research (CSIR). We thank the staff of the NCBS Central Imaging and Flow Cytometry Facility (CIFF) for help with image acquisition. Animal work in the NCBS/inStem Animal Care and Resource Center was partially supported by the National Mouse Research Resource (NaMoR) grant # DBT/PR5981 /MED/31/181/2012;2013-2016 & 102/IFD/SAN/5003/2017-2018 from the Indian Department of Biotechnology

## Author Contributions

Conceptualization, T.B. and C.J.; Methodology, T.B., C.J.; Investigation, T.B., R.D., A.M.H., A.A.K., A.J.P., A.P.D; Writing – Original Draft, T.B., C.J.; Writing – Review & Editing, T.B., C.J.; Funding Acquisition, C.J. and S.R.; Resources, C.J., and S.R.; Supervision, T.B., C.J., S.R.

## REFERENCES

1. T. J. Shaw, P. Martin, Wound repair: A showcase for cell plasticity and migration. Curr. Opin. Cell Biol. 42, 29–37 (2016).

2. M. A. Troester, et al., Activation of Host Wound Responses in Breast Cancer Microenvironment. Clin. Cancer Res. 15, 7020–7028 (2009).

3. C. J. Lewis, A. N. Mardaryev, A. A. Sharov, M. Y. Fessing, V. A. Botchkarev, The Epigenetic Regulation of Wound Healing. Adv. wound care 3, 468–475 (2014).

4. L. Rinaldi, et al., Dnmt3a and Dnmt3b Associate with Enhancers to Regulate Human Epidermal Stem Cell Homeostasis (Elsevier, 2016).

5. Q. Shen, H. Jin, X. Wang, Epidermal Stem Cells and Their Epigenetic Regulation. Int. J. Mol. Sci. 14, 17861 (2013).

6. D. Orioli, E. Dellambra, Epigenetic Regulation of Skin Cells in Natural Aging and Premature Aging Diseases. Cells 7, 268 (2018).

7. L. Cooper, C. Johnson, F. Burslem, P. Martin, Wound healing and inflammation genes revealed by array analysis of “macrophageless” PU.1 null mice. Genome Biol. (2005) https://doi.org/10.1186/gb-2004-6-1-r5.

8. T. Shaw, P. Martin, Epigenetic reprogramming during wound healing: loss of polycomb-mediated silencing may enable upregulation of repair genes. EMBO Rep. 10, 881–6 (2009).

9. A. Totaro, T. Panciera, S. Piccolo, YAP/TAZ upstream signals and downstream responses. Nat. Cell Biol. 20, 888 (2018).

10. R. Yang, et al., Epidermal stem cells in wound healing and their clinical applications. Stem Cell Res. Ther. 2019 101 10, 1–14 (2019).

11. M. Senoo, Epidermal Stem Cells in Homeostasis and Wound Repair of the Skin. Adv. Wound Care 2, 273 (2013).

12. P. Lee, et al., Dynamic expression of epidermal caspase 8 simulates a wound healing response. Nature 458, 519–23 (2009).

13. P. Lee, et al., Stimulation of hair follicle stem cell proliferation through an IL-1 dependent activation of γδT-cells. Elife 6 (2017).

14. S. Ghosh, et al., Extracellular caspase-1 regulates hair follicle stem cell migration during wound-healing. bioRxiv, 548529 (2020).

15. C. Li, et al., Development of atopic dermatitis-like skin disease from the chronic loss of epidermal caspase-8. Proc. Natl. Acad. Sci. U. S. A. 107, 22249–22254 (2010).

16. T. Bhatt, et al., Sustained Secretion of the Antimicrobial Peptide S100A7 Is Dependent on the Downregulation of Caspase-8. Cell Rep. 29, 2546–2555.e4 (2019).

17. S. Fulda, Caspase-8 in cancer biology and therapy. Cancer Lett. 281, 128–133 (2009).

18. D. G. Stupack, Caspase-8 as a therapeutic target in cancer. Cancer Lett. 332, 133–40 (2013).

19. C. Liedtke, et al., Silencing of caspase-8 in murine hepatocellular carcinomas is mediated via methylation of an essential promoter element. Gastroenterology 129, 1602–15 (2005).

20. L. Rinaldi, et al., Loss of Dnmt3a and Dnmt3b does not affect epidermal homeostasis but promotes squamous transformation through PPAR-γ. Elife 6 (2017).

21. M. Lawrence, S. Daujat, R. Schneider, Lateral Thinking: How Histone Modifications Regulate Gene Expression. Trends Genet. 32, 42–56 (2016).

22. J. Du, L. M. Johnson, S. E. Jacobsen, D. J. Patel, DNA methylation pathways and their crosstalk with histone methylation. Nat. Rev. Mol. Cell Biol. 16, 519–32 (2015).

23. A. D. King, et al., Reversible Regulation of Promoter and Enhancer Histone Landscape by DNA Methylation in Mouse Embryonic Stem Cells. Cell Rep. 17, 289–302 (2016).

24. X. Guo, et al., Structural insight into autoinhibition and histone H3-induced activation of DNMT3A. Nature 517, 640–4 (2015).

25. P. Voigt, W.-W. Tee, D. Reinberg, A double take on bivalent promoters. Genes Dev. 27, 1318–38 (2013).

26. X. Trepat, et al., Physical forces during collective cell migration. Nat. Phys. 5, 426–430 (2009).

27. T. Lecuit, A. S. Yap, E-cadherin junctions as active mechanical integrators in tissue dynamics. Nat. Cell Biol. 17, 533–539 (2015).

28. D. E. Leckband, J. de Rooij, Cadherin Adhesion and Mechanotransduction. Annu. Rev. Cell Dev. Biol. 30, 291–315 (2014).

29. B. Jackson, et al., RhoA is dispensable for skin development, but crucial for contraction and directed migration of keratinocytes. Mol. Biol. Cell 22, 593–605 (2011).

30. J. Mannik, K. Alzayady, S. Ghazizadeh, Regeneration of multilineage skin epithelia by differentiated keratinocytes. J. Invest. Dermatol. 130, 388–397 (2010).

31. M. Schäfer, S. Werner, Cancer as an overhealing wound: an old hypothesis revisited. Nat. Rev. Mol. Cell Biol. 9, 628–38 (2008).

32. E. M, S. L, W. O, S. W, Promoter methylation pattern of caspase-8, P16INK4A, MGMT, TIMP-3, and E-cadherin in medulloblastoma. Pathol. Oncol. Res. 10, 17–21 (2004).

33. Y. Wu, M. Alvarez, D. J. Slamon, P. Koeffler, J. V Vadgama, Caspase 8 and maspin are downregulated in breast cancer cells due to CpG site promoter methylation. BMC Cancer 10, 32 (2010).

34. S. Cho, et al., Epigenetic methylation and expression of caspase 8 and survivin in hepatocellular carcinoma. Pathol. Int. 60, 203–211 (2010).

35. E. Hervouet, F. M. Vallette, P.-F. Cartron, Impact of the DNA methyltransferases expression on the methylation status of apoptosis-associated genes in glioblastoma multiforme. Cell Death Dis. 1, 1–9 (2010).

36. I. Pastar, et al., Epithelialization in Wound Healing: A Comprehensive Review. Adv. Wound Care 3, 445–464 (2014).

37. G. K. Patel, C. H. Wilson, K. G. Harding, A. Y. Finlay, P. E. Bowden, Numerous keratinocyte subtypes involved in wound re-epithelialization. J. Invest. Dermatol. 126, 497–502 (2006).

38. T. Bhatt, et al., Sustained Secretion of the Antimicrobial Peptide S100A7 Is Dependent on the Downregulation of Caspase-8. Cell Rep. 29, 2546–2555.e4 (2019).

39. A. Nakagawara, et al., High levels of expression and nuclear localization of interleukin-1 beta converting enzyme (ICE) and CPP32 in favorable human neuroblastomas. Cancer Res. 57, 4578–84 (1997).

40. D. Subramaniam, R. Thombre, A. Dhar, S. Anant, DNA Methyltransferases: A Novel Target for Prevention and Therapy. Front. Oncol. 4, 80 (2014).

41. D. He, et al., DNMT3A/3B overexpression might be correlated with poor patient survival, hypermethylation and low expression of ESR1/PGR in endometrioid carcinoma: An analysis of the Cancer Genome Atlas. Chin. Med. J. (Engl). 132, 161–170 (2019).

42. I. Kataoka, et al., DNMT3A overexpression is associated with aggressive behavior and enteroblastic differentiation of gastric adenocarcinoma. Ann. Diagn. Pathol. 44, 151456 (2020).

43. J.-Y. Li, et al., Synergistic Function of DNA Methyltransferases Dnmt3a and Dnmt3b in the Methylation of Oct4 and Nanog. Mol. Cell. Biol. (2007) https://doi.org/10.1128/mcb.01380-07.

44. G. A. Challen, et al., Dnmt3a and Dnmt3b have overlapping and distinct functions in hematopoietic stem cells. Cell Stem Cell (2014) https://doi.org/10.1016/j.stem.2014.06.018.

45. W. Zhang, J. Xu, DNA methyltransferases and their roles in tumorigenesis. Biomark. Res. (2017) https://doi.org/10.1186/s40364-017-0081-z.

46. K. D. Robertson, DNA methylation, methyltransferases, and cancer. Oncogene (2001) https://doi.org/10.1038/sj.onc.1204341.

47. R. E. Husni, et al., DNMT3a expression pattern and its prognostic value in lung adenocarcinoma. Lung Cancer 97, 59–65 (2016).

48. H. R. Davies, et al., Epigenetic modifiers DNMT3A and BCOR are recurrently mutated in CYLD cutaneous syndrome. Nat. Commun. 10, 1–9 (2019).

49. J. Zhang, E. Bardot, E. Ezhkova, Epigenetic regulation of skin: Focus on the Polycomb complex. Cell. Mol. Life Sci. 69, 2161–2172 (2012).

50. K.-M. Noh, et al., Engineering of a Histone-Recognition Domain in Dnmt3a Alters the Epigenetic Landscape and Phenotypic Features of Mouse ESCs. Mol. Cell (2018) https://doi.org/10.1016/j.molcel.2018.01.014.

51. D. N. Weinberg, et al., The histone mark H3K36me2 recruits DNMT3A and shapes the intergenic DNA methylation landscape. Nature 573, 281–286 (2019).

52. B. Kuehlmann, C. A. Bonham, I. Zucal, L. Prantl, G. C. Gurtner, Mechanotransduction in Wound Healing and Fibrosis. J. Clin. Med. 9, 1423 (2020).

53. S. Li, D. Yang, L. Gao, Y. Wang, Q. Peng, Epigenetic regulation and mechanobiology. Biophys. Reports 6, 33–48 (2020).

54. H. Q. Le, et al., Mechanical regulation of transcription controls Polycomb-mediated gene silencing during lineage commitment. Nat. Cell Biol. 18, 864–875 (2016).

55. M. M. Nava, et al., Heterochromatin-Driven Nuclear Softening Protects the Genome against Mechanical Stress-Induced Damage. Cell (2020) https://doi.org/10.1016/j.cell.2020.03.052.

56. Y. Ling, et al., Modification of de novo DNA methyltransferase 3a (Dnmt3a) by SUMO-1 modulates its interaction with histone deacetylases (HDACs) and its capacity to repress transcription. Nucleic Acids Res. 32, 598–610 (2004).

57. B. Li, et al., Polycomb protein Cbx4 promotes SUMO modification of de novo DNA methyltransferase Dnmt3a. Biochem. J. 405, 369–378 (2007).

58. E. Rognoni, G. Walko, The Roles of YAP/TAZ and the Hippo Pathway in Healthy and Diseased Skin. Cells 8, 411 (2019).

59. F. Li, C. A. Adase, L. J. Zhang, Isolation and culture of primary mouse keratinocytes from neonatal and adult mouse skin. J. Vis. Exp. 2017, 56027 (2017).

60. J. A. Nowak, E. Fuchs, Isolation and culture of epithelial stem cells. Methods Mol. Biol. (2009) https://doi.org/10.1007/978-1-59745-060-7_14.

61. D. D. Bikle, Z. Xie, C.-L. Tu, Calcium regulation of keratinocyte differentiation. Expert Rev. Endocrinol. Metab. 7, 461–472 (2012).

62. 62. J. Wu, J. Q. Feng, X. Wang, “In situ hybridization on mouse paraffin sections using DIG-labeled RNA probes” in Methods in Molecular Biology, (Humana Press Inc., 2019), pp. 163–171.

63. J. R. Tse, A. J. Engler, Preparation of hydrogel substrates with tunable mechanical properties. Curr. Protoc. Cell Biol. (2010) https://doi.org/10.1002/0471143030.cb1016s47.

64. S. Syed, A. Karadaghy, S. Zustiak, Simple polyacrylamide-based multiwell stiffness assay for the study of stiffness-dependent cell responses. J. Vis. Exp. 2015 (2015).

65. M. Brand, S. Rampalli, C. P. Chaturvedi, F. J. Dilworth, Analysis of epigenetic modifications of chromatin at specific gene loci by native chromatin immunoprecipitation of nucleosomes isolated using hydroxyapatite chromatography. Nat. Protoc. (2008) https://doi.org/10.1038/nprot.2008.8.

66. S. A. Miller, D. D. Dykes, H. F. Polesky, A simple salting out procedure for extracting DNA from human nucleated cells. Nucleic Acids Res. 16, 1215 (1988).

67. A. M. Bolger, M. Lohse, B. Usadel, Trimmomatic: A flexible trimmer for Illumina sequence data. Bioinformatics 30, 2114–2120 (2014).

68. D. Kim, B. Langmead, S. L. Salzberg, HISAT: A fast spliced aligner with low memory requirements. Nat. Methods 12, 357–360 (2015).

69. P. D. Thomas, et al., PANTHER: A library of protein families and subfamilies indexed by function. Genome Res. (2003) https://doi.org/10.1101/gr.772403.

